# Myosin II controls junction fluctuations to guide epithelial tissue ordering

**DOI:** 10.1101/078204

**Authors:** Scott Curran, Charlotte Strandkvist, Jasper Bathmann, Marc de Gennes, Alexandre Kabla, Guillaume Salbreux, Buzz Baum

**Affiliations:** MRC Laboratory of Molecular Cell Biology, UCL, Gower Street, London, WC1E 6BT, UK.; The Francis Crick Institute, 1 Midland Road, London, NW1 1AT, UK.; London Centre for Nanotechnology, University College London, UK.; Department of Mechanical Engineering, Cambridge University, UK.

## Abstract

Homophilic interactions between E-Cadherin molecules generate adhesive interfaces or junctions (AJs) that connect neighbouring cells in epithelial monolayers. These are highly dynamic structures. Under conditions of homeostasis, changes in the length of individual interfaces provide epithelia with the fluidity required to maintain tissue integrity in the face of cell division, delamination and extrinsic forces. Furthermore, when acted upon by polarized actomyosin-based forces, changes in AJ length can also drive neighbour exchange to reshape an entire tissue. Whilst the contribution of AJ remodelling to developmental morphogenesis has been subjected to intensive study, less is known about AJ dynamics in other circumstances. Here, using a combination of experiment and computational modelling, we study AJ dynamics in an epithelium that undergoes a gradual increase in packing order without concomitant large-scale changes in tissue shape or size. Under these conditions, we find that neighbour exchange events are driven by stochastic fluctuations in junction length, which are regulated at least in part by the level of junctional actomyosin. As a result of this behaviour, the steady increase in junctional actomyosin and consequent tension that accompanies development steadily reduces the rate of neighbour exchange and orders the tissue. This leads us to propose a model in which topological transitions, that underpin tissue fluidity, are either inhibited or biased by actomyosin-based forces, to drive, respectively, tissue ordering or deformation.

## Introduction

Epithelia play an important function as selective barriers that separate animal tissues from the external environment. This depends upon the presence of linear adhesive contacts, called adherens junctions (AJs), which bind neighbouring epithelial cells to one another (Harris and Tepass, 2010; van Roy and Berx, 2008; Tepass et al., 2001). Since epithelia must tolerate changes in cell packing, even during periods of homeostasis, it is important that the gain and loss of AJs occurs without compromising epithelial integrity. This requires AJs to be dynamic structures. In a monolayer epithelium, the birth and loss of AJs follows a characteristic trajectory as cells change neighbours. First, an adhesive contact connecting two neighbouring epithelial cells is lost leading to the formation of a four-way vertex. This is then resolved by the birth and elongation of a new AJ interface at 90° to the first. This simple process, often called a T1 transition, connects the two cells in the quartet that were previously separate from one another (Guillot and Lecuit, 2013; Nishimura and Takeichi, 2009; Takeichi, 2014). Such topological transitions provide epithelial monolayers with the fluidity required to preserve tissue integrity in the face of the disruptive influence of epithelial cell division (Gibson et al., 2006) and cell delamination (Levayer et al., 2016; Marinari et al., 2012); whilst also allowing packing irregularities and defects in the tissue to be resolved (Classen et al., 2005; Farhadifar et al., 2007). Moreover, when accompanied by a redistribution of cell mass, directed neighbour exchange events can be used to drive large-scale morphogenetic movements (Fristrom, 1988; Heisenberg and Bellaiche, 2013).

In many systems, the forces required to drive T1 transitions are generated by the molecular motor, non-muscle Myosin II, as it acts on AJ-associated actin filaments (Desai et al., 2013; Lecuit and Lenne, 2007). Through the action of Myosin II, the sliding of anti-parallel filaments, coupled to the AJ, creates localised mechanical tension that causes AJs to shorten, thereby triggering neighbour exchange. This has been especially well studied in the developing Drosophila germ-band, where polarized junctional actomyosin (Simoes Sde et al., 2010), actomyosin flows (Bertet et al., 2004; Rauzi et al., 2010), together with destabilisation of E-cadherin at dorsal-ventral adherens junctions (Tamada et al., 2012) drive tissue elongation (Blankenship et al., 2006; Irvine and Wieschaus, 1994; Simoes Sde et al., 2010). Nevertheless, the impact of actomyosin-based forces on individual AJs and on the tissue as a whole critically depends on the precise localization and polarity of the actomyosin network. Thus, while a pulsed polarized actomyosin network drives neighbour exchange (Zallen and Blankenship, 2008; Zallen and Wieschaus, 2004), medial actomyosin pulses tend to induce apical cell constriction, as seen during ventral furrow invagination (Martin et al., 2009; Mason et al., 2013; Vasquez et al., 2014) and dorsal closure (Solon et al., 2009).

Neighbour exchange events have also been suggested to play a much more general role in maintaining the balance between order and disorder in epithelia (Farhadifar et al., 2007; Marinari et al., 2012). However, under conditions of balanced growth or stasis, it is not yet known whether or not actomyosin plays a direct role in the process of neighbour exchange. To address this question, here we utilise the *Drosophila* pupal notum to explore the regulation and function of junction dynamics in an epithelium, during a period in which it remains relatively stable in size and shape (Bosveld et al., 2012; Guirao et al., 2015). Strikingly, in this case, the impact of Myosin-dependent tension on neighbour exchange is different from that previously described. Instead of driving neighbour exchange, Myosin II dampens junction fluctuations to limit neighbour exchange. Subsequently, a steady rise in the level of junctional Myosin II induces the gradual refinement of tissue packing; leading to the generation of a well-ordered tissue as metamorphosis reaches an end.

## Results

For this analysis, we began by examining apical junction dynamics in flies expressing endogenous levels of E-Cadherin-GFP (Huang et al., 2009). We chose to focus our analysis on cells outside of the midline to avoid studying the neighbour exchange events that accompany epithelial cell delamination (Figure 1A) (Marinari et al., 2012). To facilitate the analysis of cellular dynamics, images were taken at a high frame rate (30 sec intervals), between 12 and 13.5 h APF, prior to the onset of cell division, thereby avoiding the impact of cell rounding upon fluctuations in length of apical cell-cell contacts (Bosveld et al., 2012; Guirao et al., 2015) (Movie S1). Importantly, neighbour exchange events were observed throughout this period (Figure 1B, C, F, G and Movie S2), at a rate of 8.5 ± 1.5 ⋅ 10^−4^ T1 events per minute per junction (Figure 5J and 7D).

**Figure 1:**
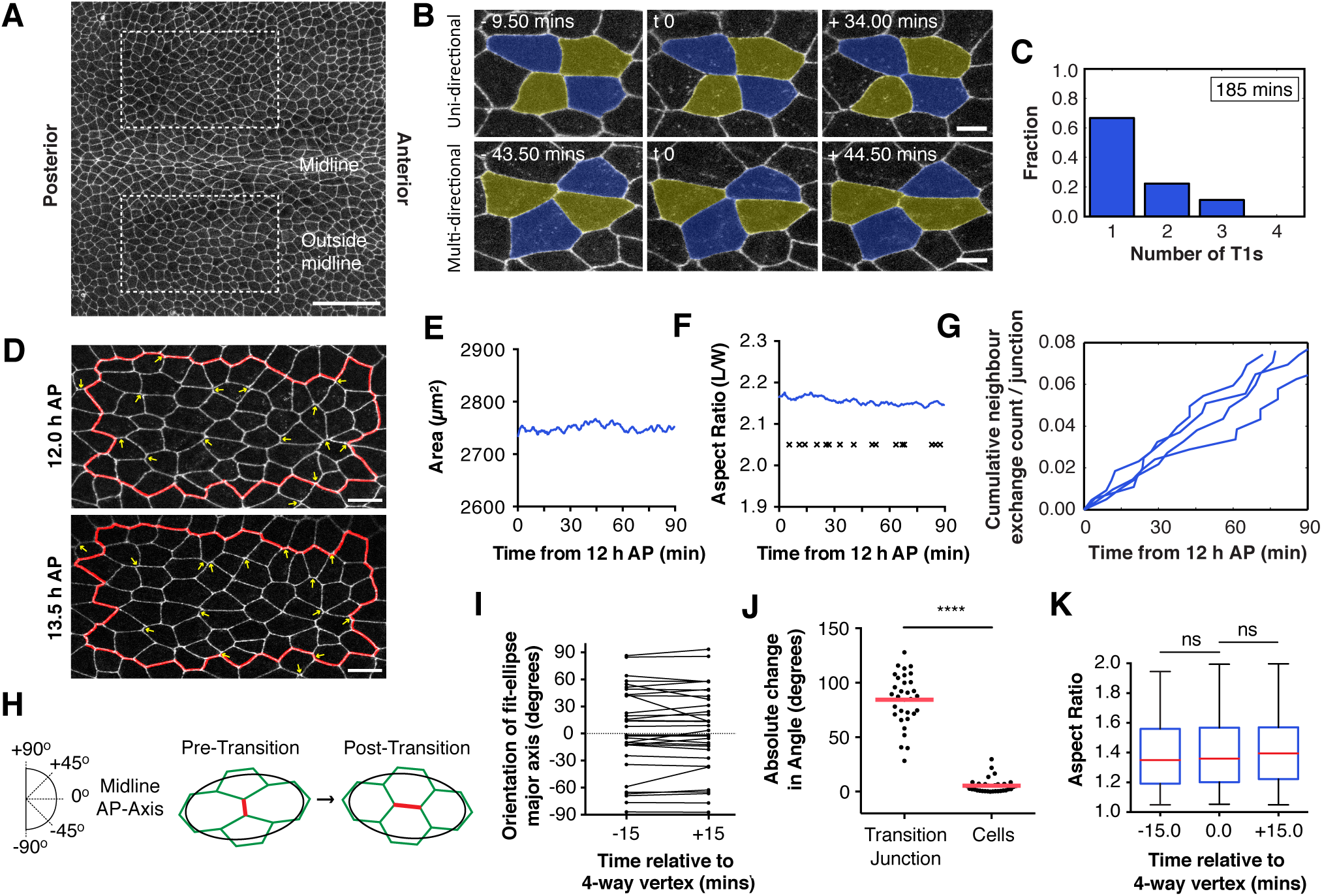
Neighbour exchange events do not contribute to tissue morphogenesis. (**A**), Apical surface projection of a live Drosophila notum labelled with DE-cadherin-GFP at 12 h AP. Image regions are indicated by dashed boxes. Scale bar, 50µm. (**B**), Neighbour exchange events are reversible. Top, Example of a uni-directional neighbour exchange event and Bottom, a multi-directional neighbour exchange event. Scale bar, 5µm. (**C**), Bar graph showing the fraction of multidirectional transition events for a representative fly. See Figure S1A for further n. (**D**), Representative region of the notum, outside the midline, at 12.0 and 13.5 h AP. Yellow arrows at 12 h label junctions that are lost, and at 13.5 h label junctions that have been gained through neighbour exchange events. Scale bar, 10µm. (**E-F**), Line plots showing the area (**E**) and aspect ratio (**F**) of the virtual clone of cells shown within the red line in D, measured over 90 mins at 30 s intervals. Crosses in **F** mark the temporal position of neighbour exchange events during this time (each cross represents the time at which a 4-way vertex is formed). (**G**), Cumulative frequency, for four individuals, of neighbour events over a 90 min period, normalised to the number of junctions within the frame at 12 h AP. (**H**), Diagram of a neighbour exchange event. The red junction represents the transition junction that is lost and gained. An ellipse (black) is fit to the four cells involved in the transition, and the feret angle of the ellipse is measured with respect to the midline (0). (**I**), The feret angle of fit ellipses are plotted at t=-15, and t=+15 mins, with paired results connected by a line. (**J**), The absolute change in angle from t=-15 to t=+15 mins for each of the four cells undergoing a neighbour exchange, and the transition junction itself (red junction in **h**) are plotted as a scatter dot plot. Red line represents the mean change in angle (transition junction = 84.28, cells = 5.342). (**K**), Box and whisker plot of the aspect ratio of the four cells undergoing neighbour exchange at-15, 0 and +15 mins relative to the formation of a four-way vertex. Red line represents median, box represents 25 and 75 percentile, tail represents the full data range. A paired t-test was used to compare t=-15 to t=0 (p = 0.7467), and t=0 to t=+15 (p = 0.1490). n for **I**, **J** and **K**= 32 exchange events from 5 flies.

One of our initial goals was to compare the junction dynamics of this system to those described for germ-band elongation (GBE) in the fly embryo (Irvine and Wieschaus, 1994). Here, cell intercalation has been proposed to proceed via relatively discrete steps that occur in sequence to make the process irreversible (Bertet et al., 2004). During GBE, dorsal-ventral oriented junctions are lost, leading to the formation of 4-way vertices. New junctions are then formed and expand in the perpendicular direction to elongate the embryo along the anterior-posterior axis (Irvine and Wieschaus, 1994). Unlike in the germ band, neighbour exchange in the notum proved to be a reversible process (Figure 1B). In many instances quartets of cells underwent multiple rounds of neighbour exchange within the imaging period (75-180 mins) (Figures 1B and C, S1A and B). Equally, many cells reaching a four-way vertex failed to undergo a neighbour exchange event, so that the ‘lost’ junction was later reformed restoring the original local tissue topology (not shown).

In contrast to the developing germ-band, where polarized neighbour exchange events drive a large-scale change in tissue shape, there was no apparent pattern to the timing or position of neighbour exchange events in the notum (Figure 1D). Neighbour exchange in the notum was not accompanied by a significant global change in tissue area (Figure 1E) or aspect (width/length) ratio – which remained nearly constant over the 90 min imaging period (Figure 1F). This was also true at the local level. There was no obvious orientation bias in AJ organisation, including no correlation between the length of AJs and their orientation in the notum (Figure S3C). Moreover, there was no bias in the orientation of junctions lost or gained during a neighbour exchange event relative to the overall tissue axis (Figure 1H and I). Most significantly, perhaps, whilst neighbour exchange in the notum involved a 90° change in the orientation of AJs, as is characteristic of T1s in other tissues, this local change in junction topology was not accompanied by a change in the shape of the associated four-cell cluster (Figure 1J and K). In other words, at the level of AJs, the exchange of neighbours at the centre of a four-cell cluster had no influence on the overall form of the cluster’s outer edge. Thus, neighbour exchange events in this tissue are reversible, do not lead to an overall tissue deformation, and occur in the complete absence of cell division, delamination and morphogenesis.

These observations led us to explore whether neighbour exchange events in the notum might arise from fluctuations in junction length, which are a ubiquitous feature of the system. To address this question we began by asking whether the length fluctuations associated with neighbour exchange were distinct from those measured in the junction population as a whole (Figure 2). Using persistence length (lengths associated with periods of ballistic motion) as a measure, we found that fluctuations around neighbour exchange events are statistically indistinguishable from fluctuations in AJ length more generally (Figure 2A-D and S3). In addition, during neighbour exchange events, the dynamics of junctional loss and gain were kinetically indistinguishable, and were no different from those measured during the shortening and lengthening of longer fluctuating junctions in the tissue (Figure 2E). These data suggest that AJs are subject to length fluctuations, a proportion of which fluctuate through zero, triggering neighbour exchange.

**Figure 2:**
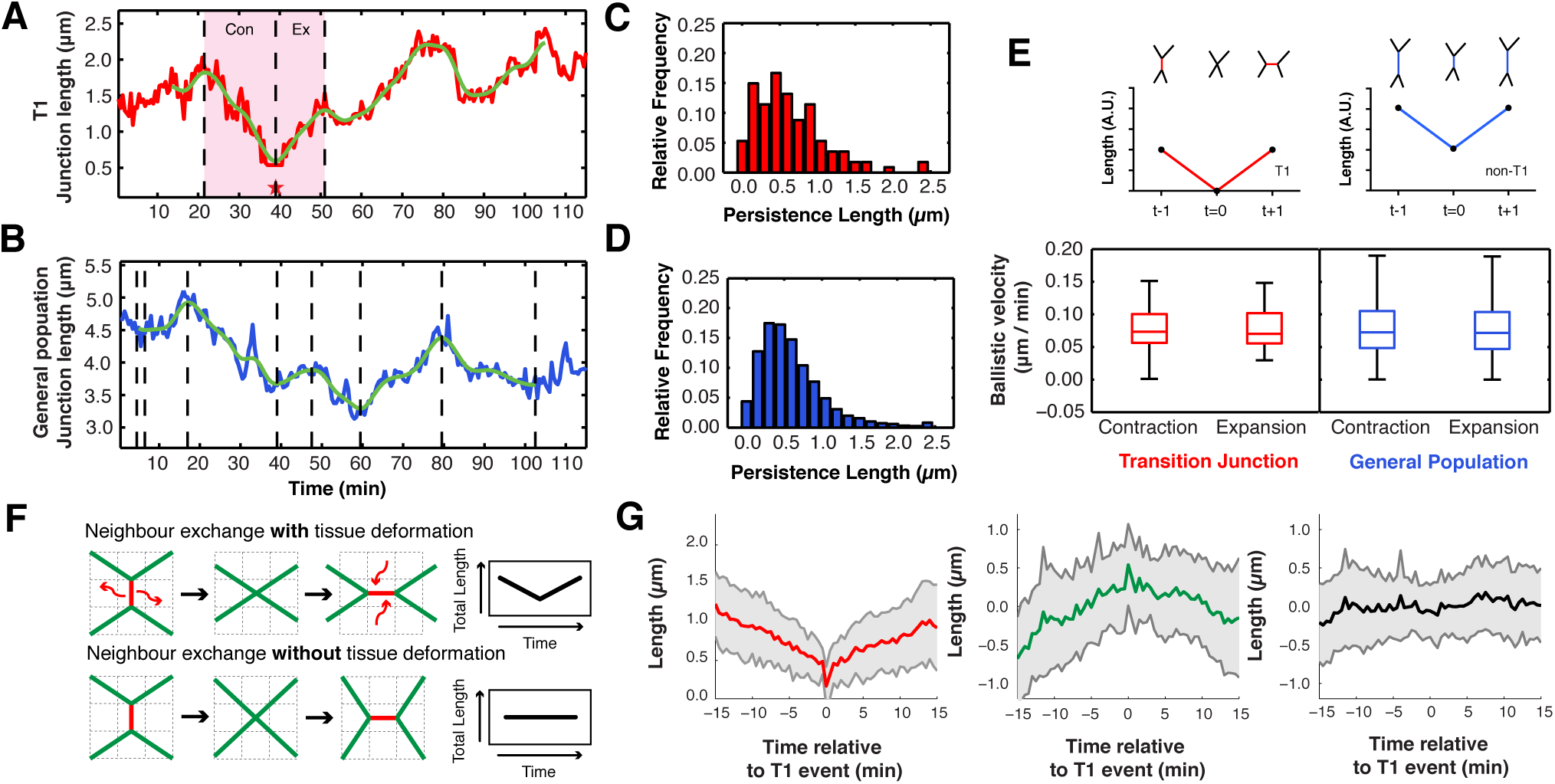
Neighbour exchange events resemble stochastic fluctuations in junction length. (**A-B**), Time series line plot for (**A**) a junction undergoing a T1 transition and (**B**) a junction fluctuating in length but not undergoing a T1. (**C-D**), Persistence length distributions for (**C**) T1 and (**D**) non-T1 junctions. The two distributions were compared using a two-sample Kolmogorov-Smirnov test, p-value = 0.5294 (ns). (**E**), (Top) Sketch of two junctions undergoing exact changes in junction length, at the same rate. Whilst the red junction undergoes a neighbour exchange, the longer junction, in blue, does not. (Bottom) Boxplots for the ballistic velocity (persistence length / persistence time) for non-T1 and T1 segments measured *in vivo*. The p-values for a Kolmogorov-Smirnov test comparing the distributions are: contracting non-T1 vs expanding non-T1 p = 0.1613 (ns), contracting T1 vs expanding T1 p = 0.9594 (ns), all non-T1 vs all T1 p = 0.3243 (ns). The filter setting is 20. N = 11377 non-T1 segments and 114 T1 segments from 4 nota. (**F**), Alternative models of neighbour exchange. Top, Developed from germ band elongation. A junction is actively removed during loss and added during expansion. This causes total junction length to decrease and then increase. Bottom, Junction material and total junction length is maintained throughout a transition. This requires slight movement of first neighbour vertices. (**G**), Average junction length changes during T1 events for wild type tissue. Plots are colour-coded according to the junction colours in **F**. (Left, Red) Line plot of the T1 transition junction, (Middle, Green) First neighbours to T1 junction, (Right, Black) Cumulation of all five junctions involved in the transition. Grey, indicates S.D. N = 27 events, from 4 nota. For each event, the junction length time series has been aligned to the four-way vertex configuration occurring at t = 0.

Previous work on GBE has suggested that neighbour exchange involves discrete steps: shrinking junctions are first lost, generating a 4-way vertex, before a new junction can be formed (Figure 2F). Previous studies have shown that these local neighbour exchange events take ~15 mins (Bertet et al., 2004; Kasza et al., 2014). While neighbour exchange events in the notum followed a similar path, the process was much slower than in the embryo. During a germ band T1 transition, one expects to observe a transient reduction in total junction length, as a junction is lost to generate a four-way vertex, followed by an increase in total junction length during the expansion phase (Bertet et al., 2004). Strikingly, this was not what we observed in the notum. Here, the extent of junctional material loss was matched by the length added during the subsequent expansion. This ensured that, on average, total junction length remained constant over time (Figure 2G). Furthermore, the process was smooth (Figure 2G, right; although a small subset of junctions paused at the four-way vertex (data not shown) so that the extent of average junctional loss/gain was offset by a corresponding increase/decrease in the length of its four first neighbours.

### Roles for non-muscle Myosin II in driving fluctuations in junction length and neighbour exchange in the notum

These data strongly suggest that the process of neighbour exchange in the notum is different to that observed during GBE, where neighbour exchange is driven by polarized, pulses of actomyosin (Bertet et al., 2004; Rauzi et al., 2010); and where junction contraction is mechanistically distinct from expansion (Collinet et al., 2015). This doesn’t, however, rule out a role for actomyosin in the process. To explore the potential role of actomyosin-based forces during neighbour exchange in the notum we first labelled non-muscle Myosin II in the tissue using live imaging and fixed staining. This revealed the presence of a junctionally associated pool of Myosin II (Figure 3A). Importantly, the junctional localisation of this pool depended on the presence of adherens junctions, and was lost following RNAi-mediated silencing of beta-catenin or DE-cadherin (Figure 3A and B) – implying an association of this pool of Myosin II with the junction itself. Although the levels of this junctional pool of Myosin II varied between junctions across the tissue (being higher at shorter junctions and at vertices (Figure S2A and F)), the overall distribution of Myosin was not evidently polarized (Figure 4A); nor was it associated with junctions of particular orientations (Figure 4B). Similarly, Bazooka was not polarized (Figure S2D)), nor was it preferentially associated with junctions that had low levels of Myosin II (Figure S2E)-as reported for other tissues (Nakayama et al., 2008; Simoes Sde et al., 2010). Nevertheless, there was a correlation between junctional length and Myosin II density across the tissue (Figure 4C). We therefore analysed in more detail the time evolution of junctional Myosin II-GFP intensity and junctional length (Figure 4D). Strikingly, junctional Myosin II intensity fluctuates over time by about 10% around its mean value (Figure 5C). Furthermore, during phases when they are shorter, junctions tend to have slightly higher levels of non-muscle Myosin II per unit length than longer junctions (Figure 4D). Consistent with this, a cross-correlation analysis showed that Myosin II level is negatively correlated with junction length, and the accumulation of non-muscle Myosin II at AJs preceded junction shortening (Figure 4E). These data show that the junctional pool of Myosin II acts like a spring, generating tension to reduce the length of AJs in the notum.

**Figure 3:**
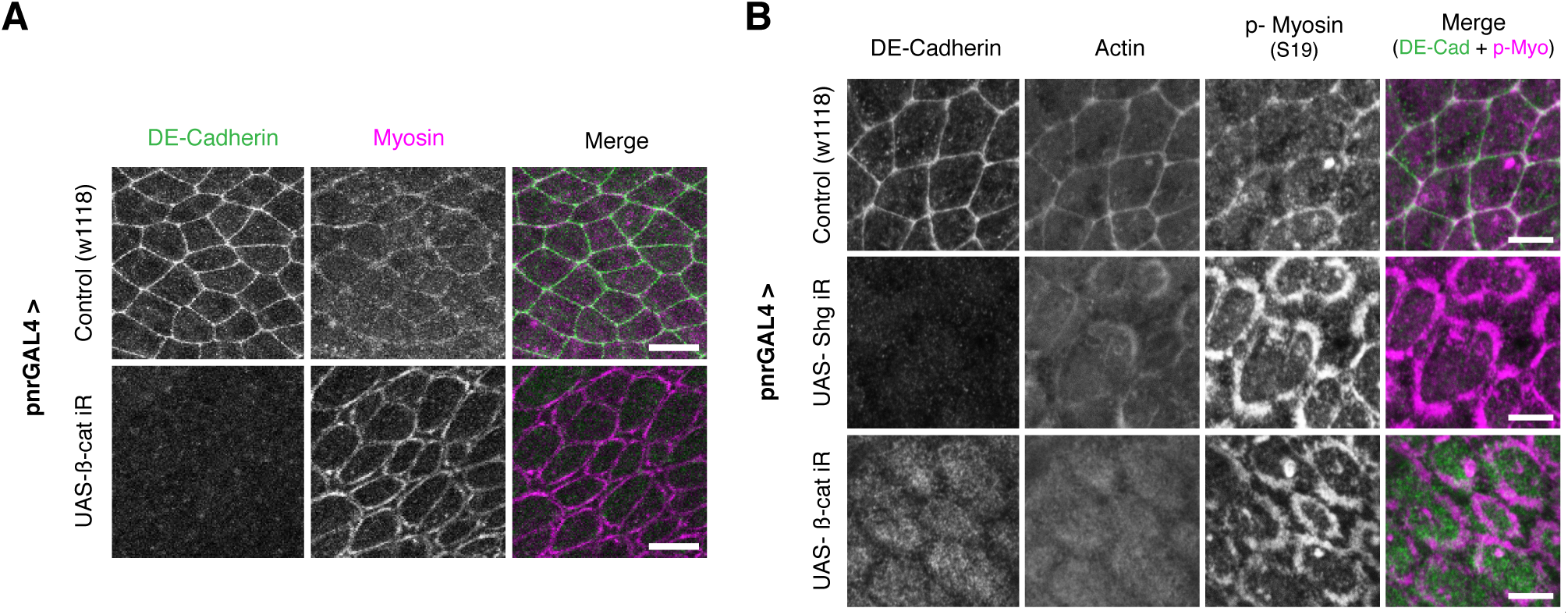
E-cadherin couples the actomyosin cytoskeleton to the apical adherens junction. (**A**), Apical surface projection from a live pupae expressing ubi-E-cadherin-GFP (Green) and MRLC-mCherry (Magenta) for Control and *UAS-β-catenin* (*armadillo*) RNAi. Scale bar, 25µm. (**B**), Representative nota of Control, UAS-*Shotgun* (*DE-cadherin*) RNAi and UAS-*β-catenin* RNAi, driven by pnr-GAL4. Tissues were fixed and stained for E-cadherin (anti-GFP against DE-cadherin-GFP), F-actin (Phalloidin) and phospho-Myosin II (S19). Scale bar, 5µm.

**Figure 4:**
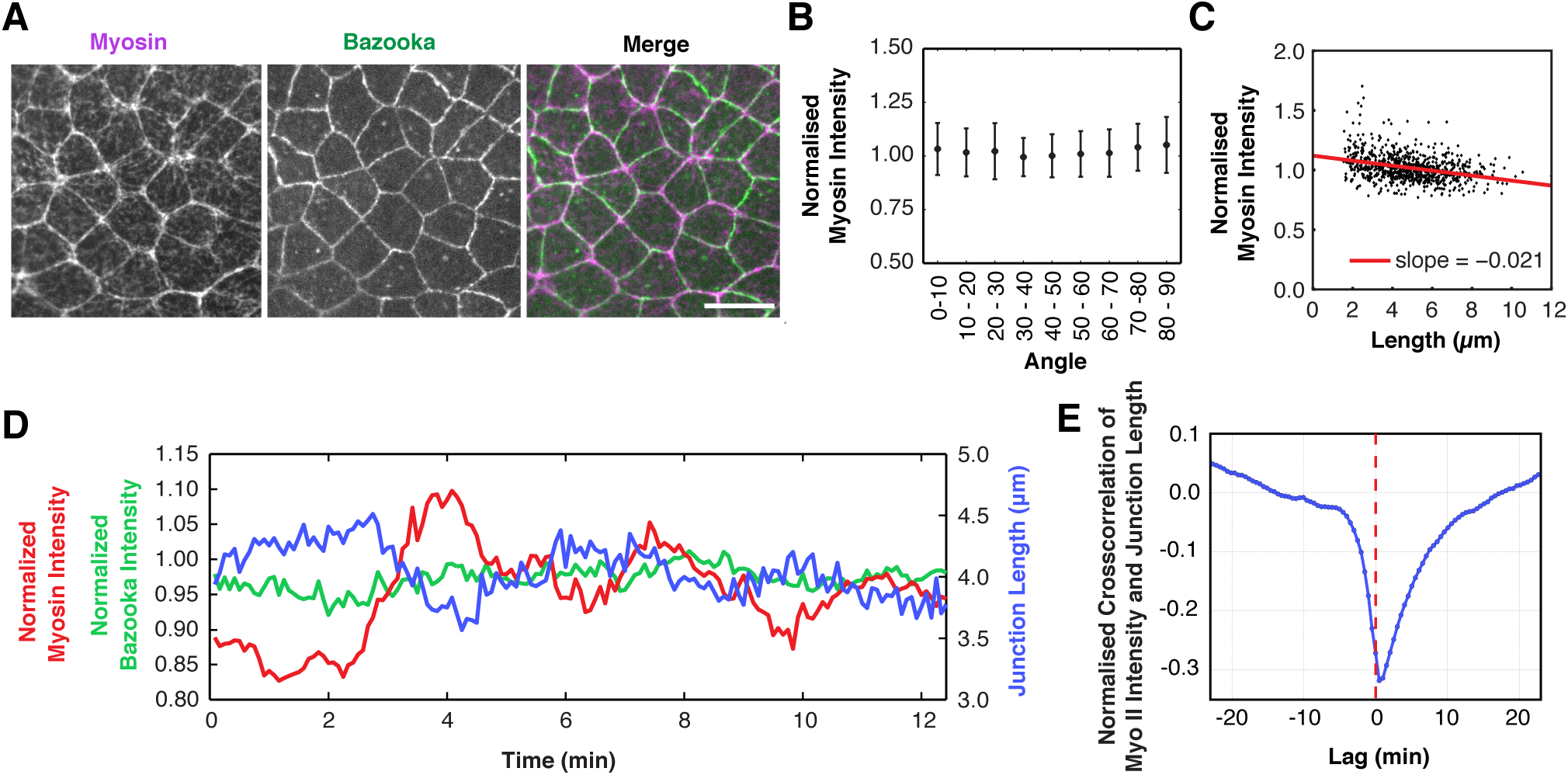
Myosin II drives changes in junctional length. (**A**), Apical surface projection of a live nota imaged with Spaghetti-Squash-GFP (Myosin regulatory light chain) (Magenta) and Bazooka-mCherry (Green). Scale bar = 5μm. (**B**), Myosin II intensity (normalised to mean intensity in a single frame at 12 h AP) versus angle, with respect to the AP midline (0). (**C**), Normalised Myosin-GFP intensity of individual junctions versus junction length. (Slope =-0.021, R^2^ = 0.21. Spearman’s Rank =-0.47. n for b and c = 809 junctions / 3 nota). (**D**), A line plot showing a representative example of junction length (Blue), Myo-II-GFP intensity (Red), Baz-mCh intensity (Green) plotted as a function of time. Both Baz-Ch and Myo-GFP intensities are normalised to mean tissue intensity. (**E**), Mean normalised crosscorrelation for Myosin intensity and junction length 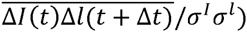 as a function of lag time Δ*t*, with σ^I^ and σ^l^ the intensity and length S.D. The normalised crosscorrelation is calculated for each junction and then averaged over different junctions (n = 269 junctions / 1 nota, imaged at 30 s intervals for 110 mins).

### Using computational modelling to determine how Myosin-dependent junction tension likely influences neighbour exchange

In order to better understand how junctional Myosin II might function at AJs to control neighbour exchange events in this monolayer epithelium, we developed a novel stochastic vertex model. In this model, forces acting on vertices arise from line tensions acting on cell-cell interfaces and from a cell area elasticity term constraining the apical cell area to a target area, as in previous studies (Farhadifar et al., 2007; Marinari et al., 2012). We also assume that vertices are subjected to a dynamic friction force, such that the equation of motion of the position of a vertex *x*_*i*_ is given by:

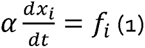

with *α* a friction coefficient, and the force *f*_*i*_ dependent in particular on the line tensions along cell bonds, *y*_*ij*_ (Figure 5A). To account for the fluctuations in junction length observed *in vivo*, we also introduced stochastic fluctuations in line tensions into the model (Equation 2). Under this simple assumption, the dynamic evolution of each vertex now depends both on the fluctuating forces to which they are subjected and on a friction coefficient that determines how quickly they respond to external forces.

**Figure 5:**
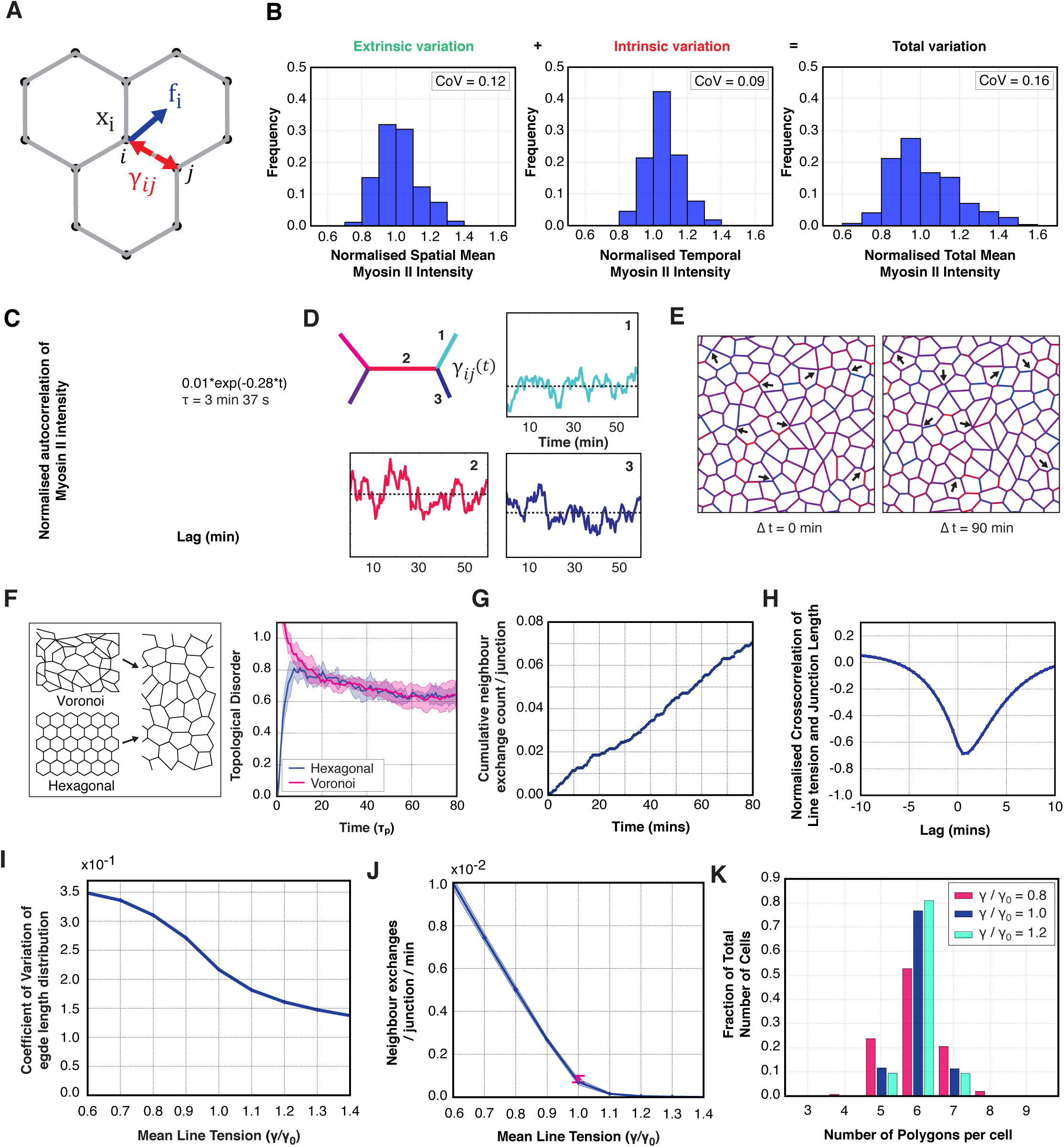
A 2D vertex model, with fluctuating line tensions, recapitulates experimental rates of topological transitions. (**A**), Schematic of the vertex model: each vertex i is subjected to a mechanical force *f*_*i*_, that depends on line tensions across cellular interfaces 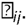. (**B**), Left: Histogram for the distribution of time-averaged junctional Myosin II intensities, for different junctions in the tissue (extrinsic variation) (n= 269 / 1 fly, CoV = 0.12, 110 mins = 220 time points); Middle: Histogram for the distribution of Myosin intensities over time for one representative junction (intrinsic variation) (n = 220 time points = 110 mins, CoV for individual junction = 0.09, full distribution of CoVs in Figure S3H); Right: Histogram showing the spatial distribution of Myosin II intensities for a single timeframe at 12 h AP (total variation). (n= 269 junctions / 1 fly). (**C**), Autocorrelation for Myosin II intensity variation on single junctions 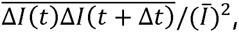, with 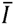 the mean intensity of the junction, as a function of lag time ∆*t* (blue). Red curve: best fit for an exponential of the form *y* + *a* * *exp* (− *b* * *x*). The coefficients are: *a* = 0.0103 ± 0.00017, *b* =-0.288 ± 0.0077. n = 269 junctions / 1 nota. (**D**), In the stochastic vertex model, line tensions fluctuate over time according to an Orstein-Uhlenbeck process, with mean tension varying over different edges. 1-3 are realizations of line tensions over time. The mean of each junction’s fluctuations 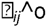 (dotted lines) are taken from a normal distribution with mean γ. (**E**), Representative example of one simulation at two time points. The edge colour, for each interface, corresponds to the level of line tension relative to its mean (blue = low, red = high). Neighbour exchange events occur when an edge length falls below a threshold. Black arrows at ∆*t* = 0 min label example junctions that are lost, and at ∆*t* = 90 min label junctions that have been gained through neighbour exchange. (**F**), Topological disorder as a function of time for two different initial packing configurations, regular honeycomb packing and packing obtained by Voronoi tessellation, with identical parameters. Topological disorder is defined as the standard deviation of the number of edges per cell within the tissue at one time point. Both tissues relax to the fluctuating state within a characteristic time of 20τ_*p*_. The convergence of simulations to the same average disorder indicates that the steady state of the fluctuating vertex model is independent of the initial packing geometries. (**G**) Cumulative count of neighbour exchange events for a representative simulation, with wild type parameter fitting, over a period of 80 minutes, normalised to the number of junctions in the frame at t= 0 min. (**H**), Cross-correlation of simulated line tensions and edge lengths 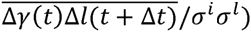 as a function of lag time ∆*t*, with σ_*i*_ and σ_*l*_ the intrinsic line tension and length S.D. (compare with Fig 4E). The normalised crosscorrelation is calculated for each junction and then averaged over different junctions. A change in line tension precedes changes in junction length by approximately 30 s. (**I**), Coefficient of variation of edge length distribution over space *CV*_*l*_, for tissues simulated with varying average line tension. The model suggests that the variation of edge lengths over different edges decreases with increasing mean line tension (γ). 1.0 represents the wild type, as measured between 12 and 13.5 h AP. (**J**), Simulated neighbour exchange rate as a function of mean line tension. Simulations reproduce the measured rate of T1 transitions for the wild type tissue (y = 1.0). The rate of neighbour exchange events is dependent on mean line tension. Higher line tensions lead to a decrease in the T1 transition rate. (**K**), Fraction of cells according to the number of neighbours. An increase in mean line tension (γ) (above 1.0), keeping other parameters constant, leads to an increase in the relative proportion of hexagonal cells.

Fluctuations in force in the system were implemented so as to mirror observed changes in Myosin II levels at individual junctions (measured using Squash-GFP; Figure 4A). In introducing these terms we were careful to distinguish between different sources of variation in levels of active Myosin II. As a measure of **extrinsic** variation, we characterized the variation in junctional Myosin II density across different junctions in the tissue by averaging the Myosin II intensity for each bond over the course of the 1.5-2 hours of observation. The resulting averages followed a near Gaussian distribution with a coefficient of variation *CV*_*e*_ ≃ 0.12 (Figure 5B, Left), showing that Myosin II intensities along separate cellular interfaces are statistically different from each other, even in the absence of tissue-wide polarised distribution. To quantify fluctuations that are intrinsic to each junction, we then quantified how the intensity of Myosin II varies at each cell-cell contact over time (Figure 5B, Middle). Intrinsic Myosin II intensity fluctuations were correlated over a period of about 3.5 minutes (Figure 5C), and the average coefficient of variation was *CV*_*i*_ ≃ 0.10; similar to the measured extrinsic variation (Figure 5B, Left). We find that the variation of Myosin II intensity across edges in the tissue at a given time, which results from both intrinsic and extrinsic fluctuations is 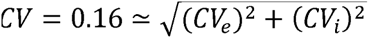 (Figure 5B, Right).

With this information in hand, we formulated a vertex model in which line tensions fluctuate over time as a result of fluctuations in Myosin II-like those observed at individual cell-cell contacts

(Figure 5D and S4). We implemented the following time-varying line tension γ_*ij*_ on the edge joining vertices *i* and *j*:

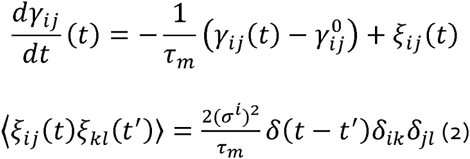

where ξ_*ij*_ is a white, uncorrelated noise characterizing intrinsic fluctuations, τ_*m*_ is a persistence time, and 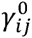 is a reference tension. In order to reflect extrinsic fluctuations, 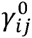 is taken from a normal distribution across edges, with mean γ and standard deviation σ^e^. We then simulated realizations of the stochastic vertex model described by equations (1) and (2) (Figure 5E). When, as a result of fluctuations in line tensions, the length of cellular interfaces falls below a defined threshold length, a T1 topological transition is induced.

Using these simulations, we first asked how cell packing is affected by initial conditions. To test this, we ran simulations starting either from a Voronoi tessellation of randomly distributed points in the plane or from a regular honeycomb packing of hexagons (Figure 5F, Left). This analysis showed that the tissue rapidly reaches a fluctuating steady state that is independent of the initial packing geometry (Figure 5F, Right). This is accompanied by a constant rate of neighbour exchange over time, as was observed in the notum (Figure 1G and S4J). In order to ensure that the parameters used in this fluctuating vertex model are close to those observed in experiments, we used the measured variations in Myosin II intensity in the tissue to set the ratio of extrinsic and intrinsic fluctuations in line tensions σ^i^/σ^e^ and persistence time τ_*m*_. We then adjusted the magnitude of fluctuations, the area elastic and the characteristic packing time, τ_*p*_ + α/γ, with *l* a characteristic cell length. For τ_*p*_ ≃ 2 minutes, line tension area elasticity ratio 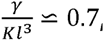 and σ^e^ ≃ 0.13γ, the fluctuating vertex model simultaneously accounts for the strength of edge fluctuations as well as the observed rate of T1 transitions (Figure 5G and S4A-I, parameters are reported in Table 1 in Model Supplement). Moreover, the cross-correlation between edge lengths and line tension fluctuations as well as the distribution of edge lengths simulated in the model are in reasonably good agreement for the model and the corresponding experimental measurements (Figure 5I and S4J-K) (Note, the distribution of junctions in the animal is skewed towards favouring short junctions, perhaps due to a subset of junctions pausing at the four-way vertex (Figure S4J)). Overall, this analysis supports the idea that T1 transitions in the tissue can be largely explained by the stochastic changes in the length of cell-cell contacts that result from intrinsic and extrinsic fluctuations in Myosin-dependent tension across AJs.

Having established a model based on the wild type tissue at 12-13.5h APF, we then wanted to determine the influence of the mean line tension on tissue dynamics. To do so we altered the mean line tension γ, while keeping other parameters constant. Under these conditions, we observed a steady decrease in length fluctuations with increasing active Myosin II (Figure 5I and Movie S3). We then used simulations to determine how neighbour exchange frequencies and topological order changed with average line tension. An increasing line tension led both to a decrease in the number of T1 transitions (Figure 5J) and to a more ordered tissue – as measured by a larger fraction of cells with 6 sides (Figure 5K). Thus, the 2D vertex model suggests that an increase in junctional Myosin II and the corresponding increase in line tension will tend to drive the tissue towards an ordered, hexagonally packed state.

### Consequences of developmental changes in Myosin II organisation

Since levels of junction tension change with developmental time in this tissue (Bosveld et al., 2012; Marinari et al., 2012), we wanted to determine if the relationship seen in the model between increasing line tension and increasing rate of topological transitions was born out during the course of notum development. To explore this question in detail, we imaged the notum over a much longer period from 20h APF. This starting point was chosen to exclude the period in which disorder is maintained by a global wave of cell division (14-20h APF). Moreover, from 20h APF, development in the tissue is accompanied by a gradual shift in Myosin II localisation, as apico-medial Myosin is lost and prominent junctional actomyosin cables are formed that appear coupled around the apical perimeter of each cell (Figure 6A). Using the junctional recoil induced by laser ablation as a measure of tension across a junction, we were able to confirm that this visible rise in the level of junctional Myosin II is associated with a significant increase in line tension (Figure 6B and Movie S4), and as suggested by previous studies (Guirao et al., 2015; Marinari et al., 2012).

**Figure 6:**
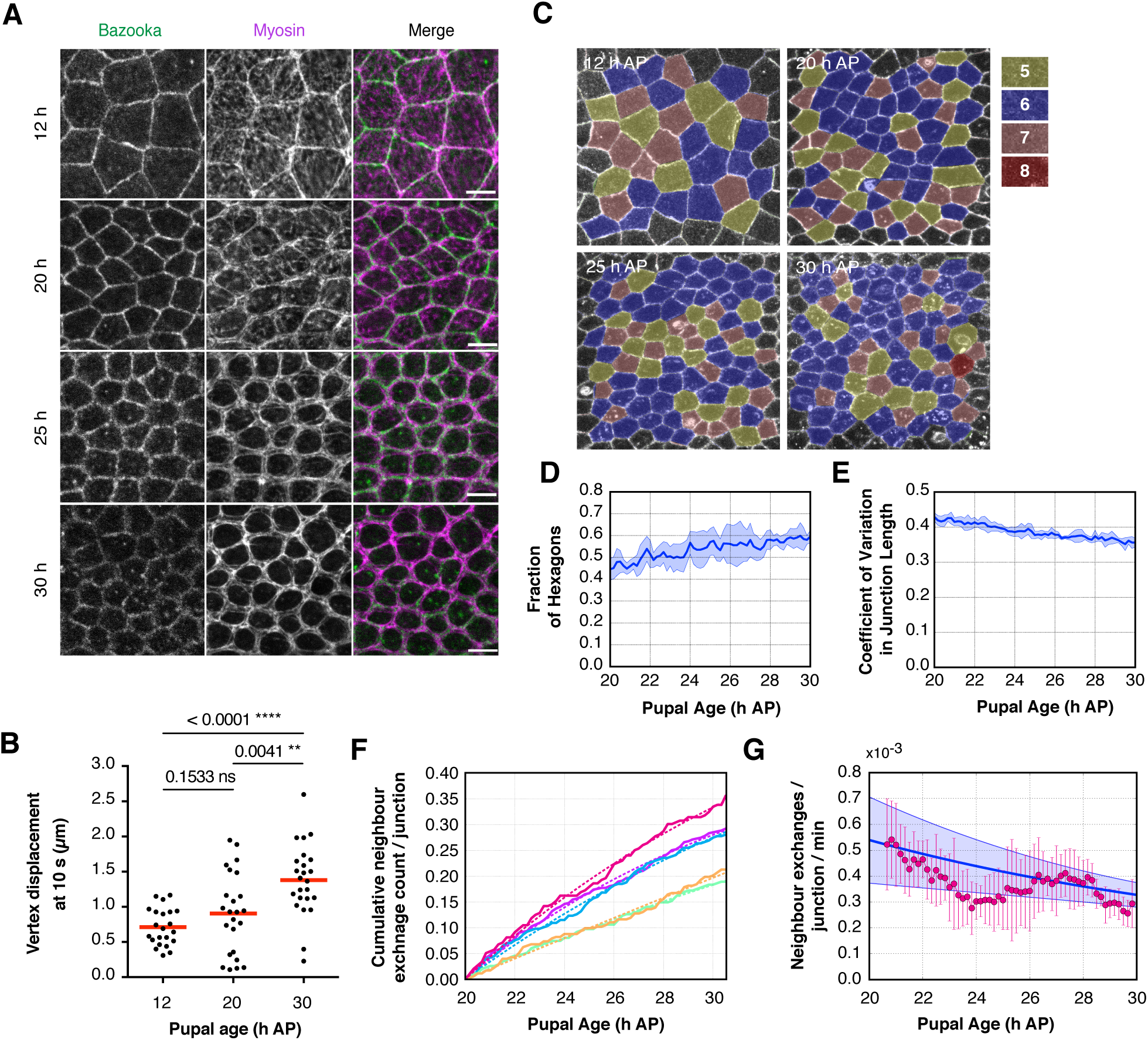
An increase in junctional Myosin II over developmental time improves tissue packing. (**A**), Apical surface projection from a live pupa expressing MRLC-GFP (Magenta) and Baz-mCh (Green). Scale bar, 10µm. (**B**), Scatter box plot quantification of the total vertex displacement at 10s of single junctions after laser dissection at 12, 20 and 30 h AP. Line indicates median. n = 22-24 cuts from 5-7 flies. (**C**), Representative regions of nota at increasing pupal development ages. Cells are coloured by number of sides. (**D**), Line graph showing the average fraction (with S.D) of hexagons within regions of nota, over time from 20 h AP. n = 86-176 cells / 5 nota. (**E**), Average (and S.D.) junction length coefficient of variation over time from 20 h APF, calculated across junctions in the tissue. n = 296 – 576 junctions / 5 nota. (**F**), Individual cumulative neighbour exchange events for each tissue, normalised to the number of junctions at 20 h APF. Dotted lines indicate a fit of the exponential (*a* *(1 − *exp*(−*b* * *t* to each individual curve. (**G**), Neighbour exchange rates from 20 to 30 h APF, inferred from **F**. Pink, T1 transition rates, with S.D., for 90 min intervals calculated from the form *T*1(*t*) + (*C*(*t* + ∆*t*/2−*C*(*t*− ∆ *t*/2))/∆*t*. Blue, mean of the derivatives (with S.D.) of the exponential fits from F.

To test whether this increase in tension is accompanied by changes in tissue order, we examined tissue packing in flies imaged from 20h through to 30h APF. As seen in the model, the observed increase in the level of junctional Myosin over time was accompanied by a significant (~15%) increase in the proportion of hexagonally packed cells (Figure 6C and D) and by a reduction in the variance of junction lengths across the tissue (Figure 6E). Furthermore, increasing line tensions were associated with an overall reduction in the rate of neighbour exchange from 20 to 30h AP (Figure 6F and G). We believe that the irregularities in neighbour exchange rate from 24-28 h are due to the growth of SOP cells, which drive local neighbour exchange (Figure S5). Thus, while the increase in junctional Myosin II is associated with a decrease in the overall rate of neighbour exchange events, the T1 transitions that do occur aid the approach to optimal cell packing.

Finally, we wanted to test whether the observed increase in the level of junctional non-muscle Myosin II across the tissue was sufficient to explain observed changes in the level of neighbour exchange events that were associated with developmental progression. Building upon previously published work, we did this by perturbing the function of the Myosin activator Rho kinase (Rok) (Simoes Sde et al., 2010; Verdier et al., 2006). We were able to confirm that Rok is present at AJs, and at four-way vertices (Figure 7A), is required for Myosin II activation (using Rok RNAi, Figure 7B), is sufficient to increase levels of junctional Myosin and p-Myosin when a constitutively active form of the kinase is over-expressed (Rok^CAT^ expressed under the Pnr-GAL4 driver, Figure 7B), and can increase the tension across AJs in the tissue (Figure 7C). Having established this, we were then in a position to use Rok^CAT^ to artificially elevate junctional Myosin II in early development (12h APF). As expected, based on the model, the increase in junctional Myosin induced by the overexpression of Rok^CAT^ led to a decrease in the frequency of neighbour exchange events – mirroring the late notum; while Rok RNAi led to a corresponding increase in neighbour exchange events (Figure 7D) (Bardet et al., 2013). At the same time, elevated levels of active Rok and Myosin II decreased the reversibility of AJs, whilst a loss of Myosin increased the chances of a T1 being reversed (Figure 7E).

**Figure 7:**
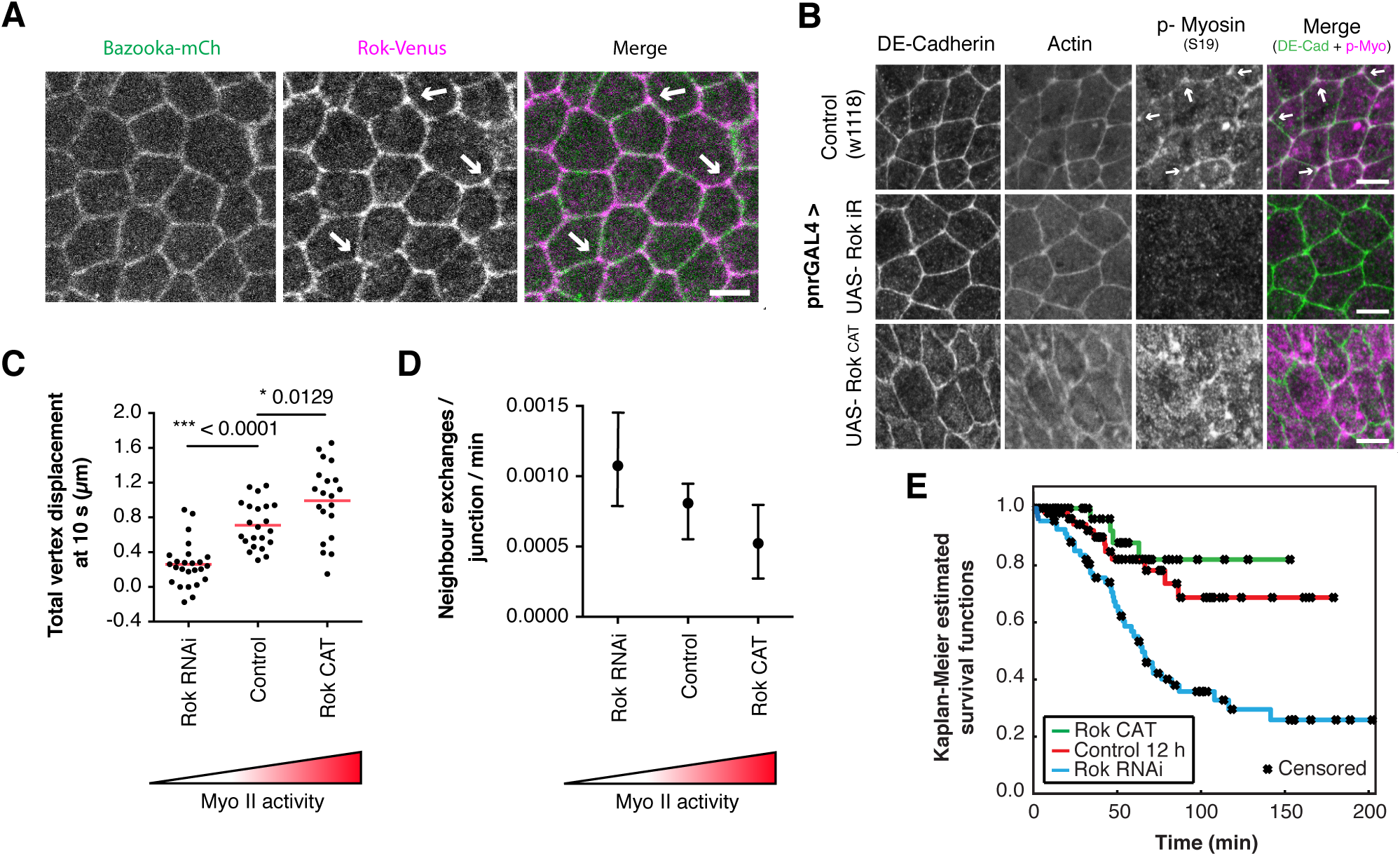
Changes in Myosin II activity tune neighbour exchange rates. (**A**), Apical surface projection of a live nota expressing Venus::Rok (K116A) (Magenta) and Baz-mCh (Green). Scale bar, 5µm. Arrows indicate enrichment of Rok at 3-way vertices. (**B**), changing levels of Myo-II through changing Rok activity are confirmed through immunohistochemistry. Arrows, in Control, indicate p-Myo II enrichment at 3-way vertices. (**C**), Quantification of total vertex displacement at 10 s after laser dissection of single junctions in 12-13.5 h AP pupae expressing Rok RNAi and RokCAT. Dots indicate individual experiments, line represents median. 4-7 flies / condition. P-values calculated from unpaired t-tests. (**D**), Quantification of normalised T1 transition rates for altered levels of Rok and subsequent Myo II activity. Dot indicates median, tails show the data range. n = 3-4 flies / condition. Mean of 232 junctions and 111 mins / experiment. Significance not shown due to low n. (**E**), Kaplan-Meier survival curves showing the probability that a neighbour exchange event is unidirectional for a given length of time. From the survival curves, the probability of a configuration persisting for at least 150 min, along with the 95% confidence interval, is: Control 0.687, [0.5175, 0.8567], UAS-Rok^CAT^, 0.258, [0.1250, 0.3915], UAS-Rok, 0.821 [0.6572 0.9841]. A log-rank test is used to determine if differences between the survival curves are statistically significant. Control vs UAS-Rok RNAi, p = 0.00041 (*), Control vs UAS-Rok^CAT^, p = 0.27399 (ns).

In sum, these data suggest that increasing the average levels of Myosin does not promote neighbour exchange in the notum – as might have been expected based upon work observed in other tissues and systems. Instead, in this relatively isotropic tissue, average junctional-associated Myosin II actively limits neighbour exchange.

## Discussion

In the context of a planar polarised tissue, the localization of Myosin II along cell-cell contacts, with a defined orientation, can generate contractile forces that contribute to tissue remodelling on a macroscopic scale, as individual cell-cell contacts are lost and new contacts form. This has been studied in detail by many groups in the context of the developing embryo. How though does Myosin-dependent tension contribute to neighbour exchange in epithelia at steady state – like the adult gut or skin?

In this paper we have explored this question by investigating the role for Myosin II in a stable epithelium, the fly pupal notum, during a period of developmental time in which there are no cell divisions, no cell delamination and no overt changes in tissue shape or size. Strikingly, in this tissue, Myosin II limits fluctuations in AJ length and, as a consequence, neighbour exchange. Therefore, with developmental time, as levels of junctional Myosin II increase, there is a corresponding decrease in the rates of neighbour exchange and an increase in tissue order, as seen by a decrease in junction length variance and by an increased proportion of hexagonal cells-both measures of improved cell packing. In this way, changes in Myosin II activity levels and localisation contribute to the refinement of the tissue observed at the end of development.

Although these observations might appear to conflict with studies of neighbour exchange in other tissues, where Myosin II has been shown to drive neighbour exchange, the function of the molecules involved seems to be identical in all cases. Thus, in the notum DE-cadherin and beta-catenin couple Myosin II to cell-cell interfaces enabling Myosin to influence tissue packing. In addition, increases in the level of Myosin II at junctions are associated with increased junction tension in the notum, just as has been described for other tissues. As a result of this, junctions with high levels of Myosin II tend to be shorter than those with low Myosin II levels in the notum. Moreover, a cross-correlation analysis shows that changes in Myosin II levels precede, and therefore likely drive, changes in junction length, as one would expect if the recruitment of Myosin II to a junction led to its contraction.

Why then is the impact of Myosin action at the level of the tissue so different in different tissues? Our model suggests that the answer may lie in the spatial organisation and temporal control of Myosin II. This is because the ability of an increase in local Myosin II at a junction to drive a neighbour exchange event depends on the state of the surrounding cell-cell junctions, which resist junction contraction. Thus, neighbour exchange will be favoured in tissues where there is a very high variance in Myosin II levels between neighbouring junctions, as exemplified by the early fly embryo, where planar polarization in the developing germ band generates extreme differences in the levels of Myosin II at perpendicular junctions (Pare et al., 2014; Simoes Sde et al., 2010), driving efficient and directed neighbour exchange. Conversely, in a tissue like the notum, where the distribution of Myosin II is relatively isotropic but fluctuates in time and space, the impact of Myosin on the rate of topological transitions will depend on the balance between the average force generated on cell bonds and the spatial and temporal fluctuations in these forces. Thus, if increased average levels of Myosin II were to induce higher levels of spatial and temporal junction length fluctuations, Myosin II would favour a more disordered tissue. Our experimental observations indicate that, in the notum, the increase in average levels of junctional Myosin and line tension seen over developmental time are accompanied by a visible decrease in the spatial variation of Myosin II (Figure 6A). As a result, the tissue becomes more ordered over time and neighbour exchange is progressively inhibited. However, our analysis suggests that epithelia can finely tune their behaviour by separately regulating average levels of junctional Myosin II, and the variation in Myosin levels between junctions across the tissue.

We note here that while the impact of Myosin II on junctional length fluctuations can explain the observed changes in tissue organisation, Myosin II may also play additional roles in the process of neighbour exchange in the tissue. This is hinted at by the observed accumulations of Myosin II and Rok at tri-cellular junctions (Figure 77 and B). Thus, in limiting neighbour exchange, Myosin II may also limit the ability of fluctuations in junction length to induce smooth passage through a 4-way vertex. This may help explain the impact of Myosin on the reversibility of junctions.

Finally, this study shows how local fluctuations in the activity, localization and levels of a molecule, in this case Myosin II, can drive local changes in cell shape, to produce larger changes in tissue topology and packing. In this way, our analysis of length fluctuations bridges the molecular, cellular and tissue scales. While this type of analysis remains in its infancy, it is likely to be important for coming to a mechanistic understanding of a wide range of biological processes. Moreover, our analysis shows how the emergence of tissue order can be driven by apparently stochastic fluctuations (Cohen et al., 2011) that are the inevitable consequence of the action of small numbers of molecules that can be tamed within a network and averaged over time, rather than by a directed developmental programme. The disadvantage of organising things in this way relate to the timescale of events. These types of noisy processes are probably too slow to be used to drive the rapid pace of morphogenesis events required by the developing fly embryo. However, organising a tissue in this way has its advantages. As a result, the epithelium will be robust to perturbations that are intrinsic, e.g. cell division and delamination, and external, such as a forced deformation. Thus, fluctuation-induced changes in cell packing, of the type we see here, would seem to be a good way of maintaining integrity in a dynamic, living epithelium. We therefore expect to see similar processes throughout the animal world.

## Experimental Procedures

Full experiments procedure can be found in the Supplementary Information.

## Acknowledgements and Funding

We are grateful to Bloomington Drosophila stock center [NIH P40OD018537], the VDRC stock center, Yohanns Bellaiche, Jennifer Zallen and Eric Wieschaus for fly stocks used in this study. We thank Paromita Majumber and Jonathan Gale (UCL) for help with the laser ablation experiments, Andrea Dimitracopoulos (MRC-LMCB, UCL) for laser ablation analysis assistance, Yanlan Mao (MRC-LMCB, UCL) for imaging assistance, Natalia Bulgakova (Univ. of Sheffield) for GBE imaging assistance & Tom Wyatt (LCN, UCL), Julia Duque (CBMSO) and Guillaume Charras (LCN, UCL) for cell culture contributions.

This work was supported by the Wellcome Trust (092840/Z/10/Z to S.C.); Cancer Research UK (C1529/A9786; C1529/A17343 to B.B.); the MRC-LMCB, University College London (S.C., C.S. & B.B.); EPSRC-CoMPLEX PhD program UCL (C.S.); the Francis Crick Institute which receives its core funding from Cancer Research UK (FC001317), the UK Medical Research Council (FC001317), and the Wellcome Trust (FC001317) (G.S., J.B. & S.C.) and ERASMUS+ (J.B.).

## Author Contributions

S.C. and B.B. wrote the manuscript. S.C. designed, performed and analysed fly experiments under the guidance of B.B. C.S. performed the junction fluctuation analysis guided by A.K. J. B. developed, designed and ran simulations in the 2D vertex model, in collaboration with M.d.G., under the guidance of G.S.

